# Overall dynamic body acceleration as an indicator of dominance in homing pigeons

**DOI:** 10.1101/2020.04.15.043349

**Authors:** Rhianna L. Ricketts, Daniel W. Sankey, Bryce P. Tidswell, Joshua Brown, Joseph F. Deegan, Steven J. Portugal

## Abstract

The benefits of dominance are well known and numerous, including first access to resources such as food, mates and nesting sites. Less well studied are the potential costs associated with being dominant. Here, the movement of two flocks of domestic homing pigeons (*Columba livia*) – measured via accelerometry loggers – was recorded over a period of two weeks. Movement was then used to calculate each individual’s daily overall dynamic body acceleration (ODBA, *G*), which can be used as a proxy for energy expenditure. The dominance hierarchy of the two flocks was determined via group-level antagonistic interactions, and demonstrated a significantly linear structure. The most dominant bird within each flock was found to move significantly more than conspecifics – on average, *c*.39% greater than the individual with the next highest degree of movement – indicating a possible cost to possessing the top rank within a hierarchy. Despite the dominance hierarchy being linear, mean daily total ODBA did not reflect a linear nature, with no pattern observed between rank and ODBA, once the top ranked individuals had been accounted for. This suggest that energy expenditure may be more reflective of a despotic hierarchy. These results show the potential for the future use of accelerometery as a tool to study the fusion of energetics and behaviour.

**Subject Category:** behaviour

**Subject Areas:** behaviour, physiology

## 1. Introduction

Group living is a common way of life for many animals [1-4]. The formation and persistence of these social groups is driven by the interests of the individuals who comprise it, not by the interests of the group as a whole [5]. For group living to persist, therefore, it must be less costly to an individual’s fitness than living alone [5]. Such benefits to fitness can be derived from decreased risk of predation [6], reduced time spent vigilant [7], improved foraging efficiency [3] and energetic savings [8]. Living in a group, however, always comes with costs which must either be tolerated or overcome [5]. These costs include increased disease transmission [4], increased detection and attack from predators [5], and increased competition for resources resulting in increased aggression [9].

One way to reduce daily aggression between members of a group is the formation of dominance hierarchies [10]. Dominance hierarchies reduce the occurrence and severity of aggressive interactions between individuals [10]. Hierarchies can be linear when dominance is established and then follow a transitive order (e.g. A>B>C and A>C) or nonlinear, when the rank order is irregular (e.g. A>B>C and C>A) [11]. These hierarchies decide the order of access to resources which are either limited, e.g. mates [10-12], with the most dominant taking the best resources. While being the most dominant individual in a group comes with clear benefits, there can also be costs associated with dominance. One such cost could be increased energy expenditure [13]; performing regular antagonistic behaviours to maintain dominance is likely to cost energy.

An individual’s basal metabolic rate (BMR) has long been assumed to influence behaviour, and a convincing argument is that a lower BMR allows higher metabolic scope to perform energy demanding activities, which may include aggressive behaviours that permit dominance [13]. Meta-analyses of multiple studies have shown that there are significant correlations between daily metabolic rate (not BMR) and traits assumed to be associated with net energy gain, such as boldness and dominance [13]; animals with higher daily metabolic rates (DMR) are more dominant, bolder, and also forage at more efficient rates [13].

Here we study two flocks of homing pigeons (*Columba livia*) to investigate the link between position within a dominance hierarchy and daily overall dynamic body acceleration (ODBA, *G*), a proxy for energy expenditure [14]. We test the hypothesis that dominant individuals within the flock will be the most active – thus most likely expending the greatest energy – to assert their dominance through antagonistic behaviours.

## 2. Material and methods

### (a) Subjects and housing

A group of 18 homing pigeons (*Columba livia*) aged 6 – 12 months old were kept in two flocks of nine pigeons each at Royal Holloway University of London (Egham UK). Flock 1 was composed of four males and five females, and flock 2 was composed of five males and four females. All pigeons had been housed together since approximately one month old in two flocks of varying composition. Sex was determined via genetic testing of feather samples. Each flock was housed in a separate loft (7ft x 6ft). The pigeons were provided with *ad libitum* access to food (Johnstone & Jeff Four Season Pigeon Corn, Gilberdyke, UK), grit and water.

### (b) Dominance Hierarchies

Determination of dominance hierarchies followed the precise protocols of [15,16]. See supplemental material for full details. The total number of interactions between individuals was recorded in a matrix, as initiators of aggressive acts (winners) or receivers of aggressive acts (losers) from each interaction [15,16]. The matrix was then used to calculate a rank for each bird using David’s Score [17], and the linearity of the hierarchy using Landau’s linearity index (h’) [18].

### (c) Overall Dynamic Body Acceleration (ODBA)

Measurement of ODBA occurred during February and March 2018. Each pigeon in both flocks was fitted with a harness which held two accelerometers (23 x 32.5 x 7.6 mm, 11g, Axivity Ltd, Newcastle upon Tyne, UK) on the centre of their backs, for a period of two-weeks. One accelerometer was programmed to record for the first week, and the other was to record for the second week to ensure full data capture while minimising disturbance. During this time all pigeons remained within their home lofts. ODBA (*G*) for each bird was calculated from the raw accelerometry data using the formula presented in [14]. Data analysis was carried out in RStudio [19,20]. An ANOVA with a Bonferroni post-hoc test was used to investigate the variation between individual’s daily ODBA in SPSS (IBM SPSS Statistics, Armonk, NY: IBM Corp.). The assumptions of parametric tests used were checked and met before tests were run.

## 3. Results

### (a) Dominance

The hierarchies of both flocks were highly linear (flock 1, h’ = 0.68, *p* = 0.006; flock 2, h’ = 0.84, *p* = <0.001). David’s score was found to correlate significantly with sex (Spearman’s rank; *rs* = 0.48, *p* = 0.04), with males being more aggressive.

### (b) Overall Dynamic Body Acceleration

Mean ODBA per hour (*G*) showed a circadian rhythmic pattern, with peaks centred around midday, and troughs throughout the night in both flocks (figure 1). An exponential decrease in total ODBA was seen with a decrease in rank (here a decrease in rank is from 1 to 10 as 1 is the highest ranked individual, and 10 the lowest) (figure 2). In both flocks, a steep decrease in total ODBA was seen within the top two ranked birds, with a percentage difference between these individuals of 45% and 32% in flock one and two respectively (figure 2). A One-Way ANOVA showed there was significant variation between individuals in both flocks (flock 1, F_8,127_ = 19.687, *p* < 0.001; flock 2, F_8,114_ = 12.567, *p* < 0.001). A Bonferroni post-hoc test showed that the most active bird in each flock was significantly different from all other birds (*p* < 0.01 for all comparisons). This was also confirmed by a Tukey HSD post-hoc test, which placed the most active bird in each flock in their own homogenous subset, indicating they had no similarity to any other bird (n = 1, *p* = 1), while all other members of the flock were found to be homogenous to at least 3 other birds.

**Figure 1.**
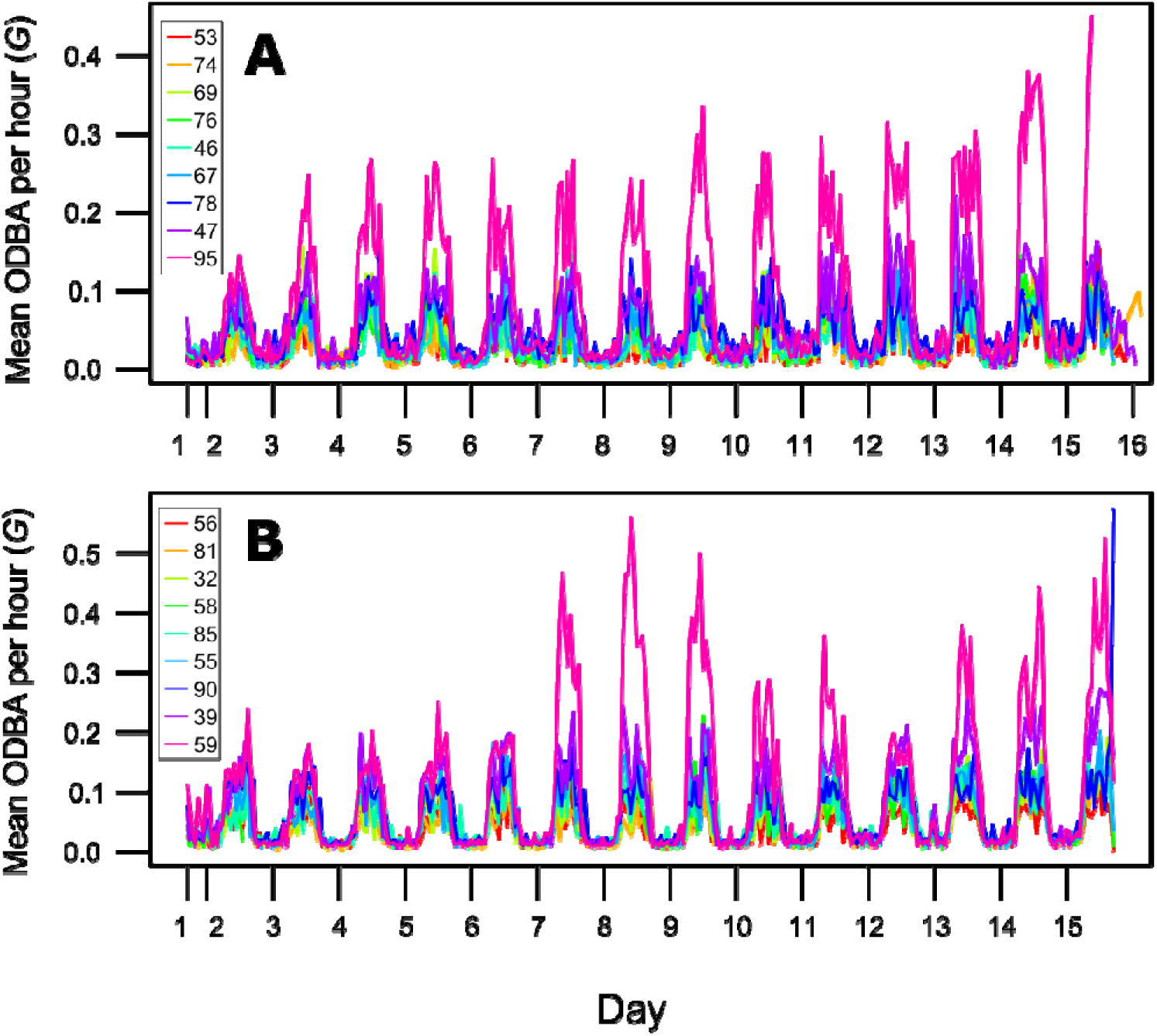
Mean ODBA per hour (gravitational constant, *G*) of nine homing pigeons (flock 1) over a two-week recording period. The x-axis tick marks indicate 5 am and 5 pm of each day, respectively. The cyan line is the number 1 ranked bird in the dominance hierarchy.

**Figure 2.**
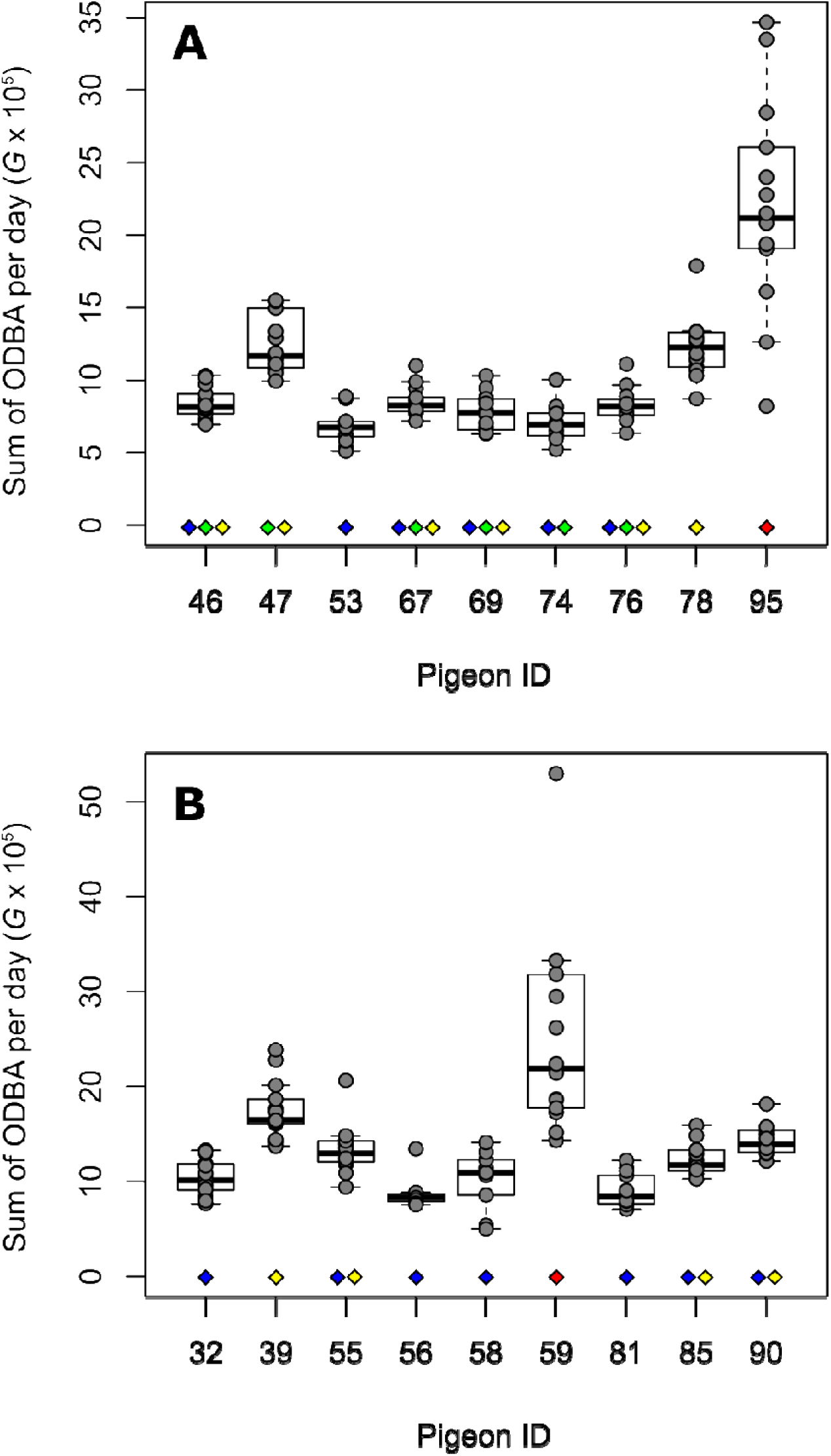
The sum of ODBA (gravitational constant, *G*) for each complete day of the recording period for all pigeons in *a*) flock 1 and *b*) flock 2. Filled diamonds indicate the homogenous subsets calculated with a Tukey HSD post-hoc test. a) Blue; subset 1, n = 6, *p* = 0.97, green; subset 2, n = 6, *p =* 0.069, yellow; subset 3, n = 6, *p* = 0.082, red; subset 4, n = 1, *p* = 1. b) Blue; subset 1, n = 7, *p* = 0.197, yellow; subset 2, n = 4, *p* = 0.302, red; subset 3, n = 1, *p* = 1.

## 4. Discussion

Using biologging technology, this study has demonstrated the potential link between dominance and the degree to which a bird moves. Within the last decade, ODBA has been put forward as a proxy for energy expenditure [14,21]. It had not yet, however, been used for fine-scale continuous recording of movement over an extended period of time.

By examining the movement of the homing pigeons as a proxy for energy expenditure, it was found that the most dominant pigeon in each flock showed significantly higher levels of movement than its conspecifics, which all moved at similar levels which were not significantly different from one another. This would suggest that there is an energetic cost incurred in being the dominant which subordinates do not have to pay. Why the dominants are more active and what behaviour they are performing during this time though is unclear. One potential explanation for the increase in movement is that dominants may be initiating the majority of agonistic interactions. For a dominant to retain its rank, and so the benefits which come with it, the individual must continue to win all antagonistic encounters against other birds in the flock [10,12,15]. An alternate explanation for why the dominant individuals are so aggressive and active could be contradictory to the idea that low BMR permits such antagonistic behaviours; these individuals have *higher* energetic requirements. This higher energetic requirement may force them to be aggressive to ensure adequate access to food, akin to ‘lead according to need’, a theory which has previously linked to motivation and leadership in group behaviour [22].

By observing antagonistic interactions, other members of the group can gain information about which individuals they are, and are not, capable of dominating, thus reducing the number of interactions needed to maintain their place in the hierarchy [10,12,15]. This reduced number of interactions needed to maintain the hierarchy could explain why the rest of the flock showed highly homogenous levels of movement at a lower level compared to the dominant. While the social hierarchy is highly linear, the distribution of energy expenditure within both flocks is reminiscent of a despotic society [12], with one individual spending energy policing the flock, while the subordinates all move on a moderately similar level. The true cost of dominance could, therefore, be that to retain dominance and gain its benefits, dominants are largely responsible for the maintenance of the hierarchy.

During the study period, all birds were kept inside and confined to their social hierarchy; behaviours were limited to feeding, sleeping, preening and social interactions. Previously it was established that ground-based dominance hierarchies do not match that of leadership during flights [23,24]. An interesting further avenue of research would be to determine how the ODBA compares for ground-based dominant birds and flight leaders, as leaders during flights are typically having to make less adjustments to their trajectories than followers [24]. Similarly, how ODBA, flight duration and flock composition interact would provide useful insight into the energetics and compromises involved in group travel [25-27].

The results of this study show that the long-term use of accelerometers is a viable method of determining individual differences in movement, and thus energy expenditure, within groups of animals. Dominants within flocks of pigeons show higher levels of movement, suggesting they either have a larger metabolic budget [13, 28] to allow such increased movement, or take on this extra movement as a cost worth paying for continued dominance.

## Ethics

All experiment protocols were approved by the RHUL local Ethics and Welfare Committee.

## Data accessibility

Data available from the Dryad Digital Repository: http://dx.doi.org/*** [*].

## Author contributions

Conceptualisation and methodology, R.L.R, D.W.S. and S.J.P. resources, S.J.P.; data collection, R.L.R, B.P.T., J.B. and J.F.D.; analysis, R.L.R and D.W.S.; writing – original draft, R.L.R. and S.J.P; writing, reviewing and editing, R.L.R,. D.W.S., B.P.T., J.B., J.F.D. and S.J.P.

## Acknowledgements

We thank the following people for useful discussions; Pat Monaghan, Dora Biro and Rebecca Thomas. We are grateful to Lara Nouri for looking after the birds.

