## Supplementary Methods for "Overall dynamic body acceleration as an indicator of dominance in homing pigeons"

*Dominance Hierarchies*

Determination of dominance hierarchies followed that of [1,2]. To determine dominance hierarchies, birds were studied in their home lofts. Prior to each experiment, food was removed at 17:00 the day before. The next morning, the pigeons were individually identified via a back-mounted sticker, and put into a pigeon carrier within their home loft, and a single feeder placed at the opposite end of the loft on the ground. The feeder had a roof and had limited space available for feeding (eight inches). Birds were released from the basket, and behavioural recordings taken of the first 30minutes of interactions between all individuals following release from the carrier. All experiments were videoed. Interactions recorded were: pecking, chasing, beak grabbing, neck grabbing and wing slapping. The total number of interactions between individuals was recorded in a matrix, as initiators of aggressive acts (winners) or receivers of aggressive acts (losers) from each interaction. All aggressive interactions were recorded on the floor around the feeder [1,2]. The matrix was then used to calculate a rank for each bird using David’s Score [3,4], and the linearity of the hierarchy using Landau’s linearity index (h’) [5].

David’s score is a method of ranking individuals within a group based on the outcome of interactions through the formula:

DS = *w + w_2_ – l – l_2_*

where *w* is the sum of an individual’s proportion of wins and *l* is the sum of an individual’s proportion of losses, and *w­_2_* and *l_2_* are the weighted proportions of wins and losses [3,4]. The weighted *w_2_* and *l_2_* scores are calculated by multiplying each individual win (or loss) proportion by the loser’s (or winner’s) *w* score, which are then summed to produce one value. This weighting provides individuals which beat those which typically win with higher scores, and those which lose to individuals that typically lose with lower scores [3,4].

Landau’s index of linearity (*h’*) uses the interaction matrix to calculate one value which describes the linearity of the group. This value ranges from 0 to 1, where 0 indicates complete equality within the group, and 1 indicates a fully linear hierarchy. A fully linear hierarchy would be one where no individuals are equal, and David’s scores are equally distributed through their range [5].

1. Portugal SJ, Sivess L, Martin GR, Butler PJ, White CR. 2017 Perch height predicts dominance rank in birds. *Ibis.* 159, 456–462.
2. Portugal SJ, Ricketts RL, Chappell J, White CR, Shepard EL, Biro D. 2017 Boldness traits, not dominance, predict exploratory flight range and homing behaviour in homing pigeons. *Philos. Trans. R. Soc. B Biol. Sci.* 372, 20160234.
3. Gammell MP, De Vries H, Jennings DJ, Carlin CM, Hayden TJ. 2003 David’s score: A more appropriate dominance ranking method than Clutton-Brock et al.’s index. *Anim. Behav.* 66, 601–605.
4. David HA. 1987 Ranking from unbalanced paired-comparison data. *Biometrika* 74, 432–436.
5. Landau HG. 1953 On dominance relations and the structure of animal societies: III The condition for a score structure. *Bull. Math. Biophys*. 15, 143–148.
